# Investigating the use of human COVID-19 rapid assays to detect antibody and antigen in domesticated dogs (*Canis lupus familiaris*) and cats (*Felis catus*)

**DOI:** 10.64898/2026.03.04.709710

**Authors:** Lauren C. Cybulska, Sarah A. Hamer, Chloe Teasdale, Glen Johnson, Gabriel L. Hamer, Jean Grassman

## Abstract

Severe acute respiratory syndrome coronavirus-2 (SARS-CoV-2), the virus that causes coronavirus disease-2019 (COVID-19) in humans, is also known to infect animals including dogs (*Canis lupus familiaris*) and cats (*Felis catus*). This study evaluated the efficacy of human COVID-19 rapid antigen and antibody tests in dogs and cats. Nasal/oral swabs from 60 animals (32 dogs, 28 cats) and serum from 40 animals (20 dogs, 20 cats) were tested. Rapid antigen tests used on respiratory swabs showed low-to-moderate sensitivity (75% dogs, 57% cats) and moderate-to-high specificity (79% dogs, 95% cats) compared to RT-PCR. Rapid antibody tests used on serum samples demonstrated low-to-moderate sensitivity (70% dogs, 50% cats) and moderate-high specificity (60% dogs, 100% cats) compared to PRNT. While imperfect, these test kits may have some utility for field surveillance studies, particularly when species-specific rapid SARS-CoV-2 assays for dogs and cats are unavailable. These test characteristics in dogs and cats are similar to the findings from studies of the same types of tests in humans which have found an average sensitivity and specificity of common commercially available kits in the US range from 50.0-84.3% and 64.5-74.3%, respectively, when used with human samples (1,2).

## Introduction

Severe acute respiratory syndrome coronavirus-2 (SARS-CoV-2) is known to cause coronavirus disease-2019 (COVID-19) in humans as well as other species including domestic dogs (*Canis lupus familiaris*) and cats (*Felis catus*), among other animals (3). In some cases, these animals can develop symptoms similar to those seen in human COVID-19 patients (4–8), but most SARS-CoV-2-infected animals are thought to be asymptomatic (9–13). Evidence suggests that pets living in homes with COVID-positive humans are more likely to test positive, pointing to human-to-animal transmission as the likely cause (14–18). SARS-CoV-2 infections in pets are not only a risk to animal health; coronaviruses mutate easily, which represents a potential for the generation of novel viruses within non-human hosts that can then be transmitted to humans (i.e., “spillover”). Some authors have stressed the importance of continued surveillance of SARS-CoV-2 in domesticated dogs, cats, and other susceptible species, to help mitigate the threat of zoonotic (i.e., animal-to-human) and reverse-zoonotic (human-to-animal) transmission through early detection and response (19–26).

Similar to their use in humans (27–29), real-time polymerase chain reaction (RT-PCR), plaque reduction neutralization testing (PRNT) (i.e., viral neutralization), and laboratory-based enzyme-linked immunoassays (ELISAs) methods are commonly used to detect SARS-CoV-2 viral ribonucleic acid (RNA) and antibodies in dogs and cats (30–32). RT-PCR and PRNT are considered gold-standard testing methods. However, these methods can be costly and require specialized laboratory equipment and trained personnel making these tests inaccessible and impractical for widespread use, particularly in low-resource settings. This highlights the importance of identifying alternative methods of testing that are more economical and allow for rapid diagnoses and accessible surveillance techniques.

While rapid antibody and rapid antigen tests (RATs) have been developed for SARS-CoV-2 detection in humans and are affordable, no such similar tests have been developed for non-human species. In the United States (US), each assay costs as little as $9.00 and they are widely available across the country online or over the counter (33). If such tests were found to be effective when used in non-human subjects, this would be a cost-effective and portable solution to aid in widespread surveillance testing or the individual testing of animals at local veterinary clinics, animal shelters, or homes. The average sensitivity and specificity of common commercially available RATs in the US range from 50.0-84.3% and 64.5-74.3%, respectively, when used with human swab samples (1,2).

With zoonotic diseases becoming an ever-increasing threat to global health – such as avian influenza and Ebola (34,35) – it is important to consider alternative testing methods to aid in widespread surveillance of these diseases across all susceptible animal populations. It is not uncommon for researchers, veterinarians, and wildlife disease specialists to use human diagnostic tests with animal samples. For example, human rapid influenza diagnostic tests (RIDTs) have been used to detect swine influenza (H1N1 and H3N2 strains) in pigs in Southeast Asia (36). To date, there is limited information about the use of human rapid tests in animals for detecting SARS-CoV-2. Since March 2020, some authors have reported the use of human SARS-CoV-2 ELISA and PCR assays with animal samples in laboratory settings (37–39), and others have successfully conducted experiments using human rapid antigen and antibody tests with dog or cat saliva, nasal swabs, and serum samples (40–43). In this study, we investigated whether commercial rapid human assays, both antibody and antigen tests, could be used to detect SARS-CoV-2 in dogs and cats in an off-label assessment.

## Materials and methods

### Study methods

We used rapid COVID-19 tests developed for humans to test dog and cat biological samples to identify whether these assays could be used by veterinarians or public health officials in surveillance studies. Dog and cat nasal and oral swab samples collected in viral transport media (VTM), a solution used to preserve biological samples, were tested with commercially available human COVID-19 RATs (Abbott BinaxNOW® Covid-19 Antigen Card Tests, Abbott Diagnostics, Scarborough, ME, USA) (44). Dog and cat serum samples were tested with human COVID-19 rapid antibody assays (EcoTest® COVID-19 IgM/IgG Rapid Test Device, Hangzhou, China) (45). Sensitivity, specificity, positive predictive value (PPV), and negative predictive value (NPV) were estimated for the rapid assays when used with animal samples. The performance characteristics of rapid antibody and rapid antigen assays were assessed by species.

The current laboratory study was conducted in a Biosafety Level 2 (BSL-2) facility from November 2021 to March 2022. Respiratory swabs and blood samples were collected from pet dogs and cats living in COVID-positive households from 2020-2021 by a field team from Texas A&M University in College Station, Texas, USA (46). Nasal swab material was stored in VTM (CDC SOP#: DSR-052-02) (47). Oral swab material was collected from cats when nasal swab samples could not be obtained (e.g., if the sterile swab was too large for a cat’s nares).

The whole blood was spun in a centrifuge to obtain sera. Gold-standard antibody and molecular testing were performed at Hamer Laboratory prior to the current study. Antibody titers were measured in serum by PRNT, and the presence of viral RNA was measured by the molecular method of RT-PCR using VTM. This involves extracting RNA and converting it to deoxyribonucleic acid (DNA) followed by amplification by PCR. The animal sera and VTM samples were stored at −80°C prior to the initiation of the current study; freezing and thawing were kept at a minimum to preserve the quality of the samples (48). The investigator was blinded to the positivity status of the VTM and serum samples.

### Real-time polymerase chain reaction protocols

The viral RNA samples were extracted at the Texas laboratory as previously described using a MagMAX CORE Nucleic Acid Purification Kit on a 96-well Kingfisher Flex System (ThermoFisher Scientific, Waltham, MA, USA) (31). A subset of samples was tested for RNA concentration on an Epoch Microplate Spectrophotometer (BioTek, Winooski, VT, USA).

Negative controls were included in the nucleic acid purification plates (VTM) and qPCR plates (PCR-grade water) with no indication of contamination (31).

### Plaque-reduction neutralization testing protocols

At the Texas laboratory, serum was tested for SARS-CoV-2 by PRNT_90_ as previously described using Isolate USAIL1/2020, NR 52381 (49,50). The use of an early lineage of SARS-CoV-2 for the PRNT was appropriate for minimizing the occurrence of false seronegative results, given findings in humans that neutralizing antibodies mounted for recently emerged variants are capable of neutralizing ancestor lineages (50).

### Antigen testing using human COVID-19 RAT test kits with VTM solution

The anterior nasal and oral swab samples stored in VTM solution (400 µL aliquots) were used to detect the presence of SARS-CoV-2 nucleocapsid antigen with a commercially available RAT (Abbott BinaxNOW® Covid-19 Antigen Card Test) (44). A curated set of pet swabs (n=60) stored in VTM, previously subjected to RT-qPCR methods, was selected for inclusion, including 25% (8/32) dog samples and 25% (7/28) cat samples that were RT-qPCR positive. These samples were stored in −80 degree freezer for up to 6 months prior to preparing aliquots for the current study. We transferred 100 µL of VTM to the RAT device in the location where the buffer is normally administered. The buffer solution that was included in the test kit was not used. The sterile swab included in each test kit was inserted in the test card after the VTM solution was administered to the device; the sterile swab was rotated in the test device, and the test card was closed. The results were read at least 15, and no more than 30, minutes after initiating the test, per the instruction manual (44).

### Antibody testing using human COVID-19 rapid antibody test kits with sera

A curated set of pet serum samples (n=40) previously subjected to PRNT methods was selected for inclusion, were tested using EcoTest® COVID-19 IgM/IgG Rapid Test Device to detect IgG and IgM antibodies. The set included 40-50 µL aliquots of serum that had been in freezer storage for up to 6 months, with 50% (10/20) dog and 50% (10/20) cat samples being true positives. The tests were used in the same manner described in the manufacturer’s instructions with the use of animal serum samples instead of human serum (45).

### Sample size and calculations

The sample size used for the study was based on the available SARS-CoV-2 positive animal samples. A sample was considered a true positive if it was positive on RT-qPCR or PRNT. The sensitivity (proportion of all true positives that showed positive test results), specificity (proportion of all true negatives that showed negative results), positive predictive value (PPV; probability that a positive test result is truly positive), and negative predictive value (NPV; probability that a negative test result is truly negative) were calculated for all test results.

### Ethical considerations

The animal samples were collected by the Hamer Laboratory team at Texas A&M University with prior approval from the Texas A&M University Institutional Animal Care and Use Committee (IACUC), located in College Station, Texas, USA, Animal Use Protocol #IACUC 2022-0001 CA.

## Results

For dog serum samples, the rapid antibody assay was found to have 70% sensitivity and 60% specificity, the PPV was 64% and the NPV was 67% in comparison to PRNT methods (Table 1). For cat serum samples, the rapid antibody assay was found to have 50% sensitivity and 100% specificity, the PPV was 100% and the NPV was 40%. All the positive tests had confirmed IgM indicator lines and there were no IgG-positive samples. Some of the IgM test bands were very faint, while other test bands were as dark as, or darker than, the control line.

**Table 1.**
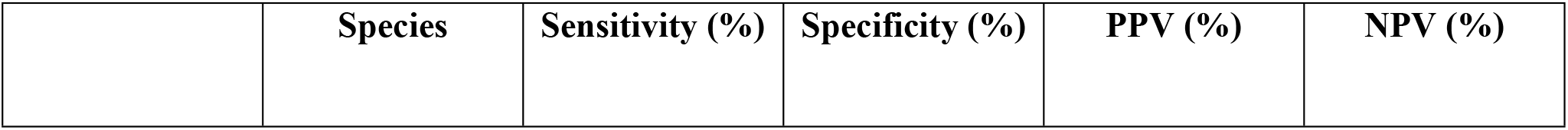
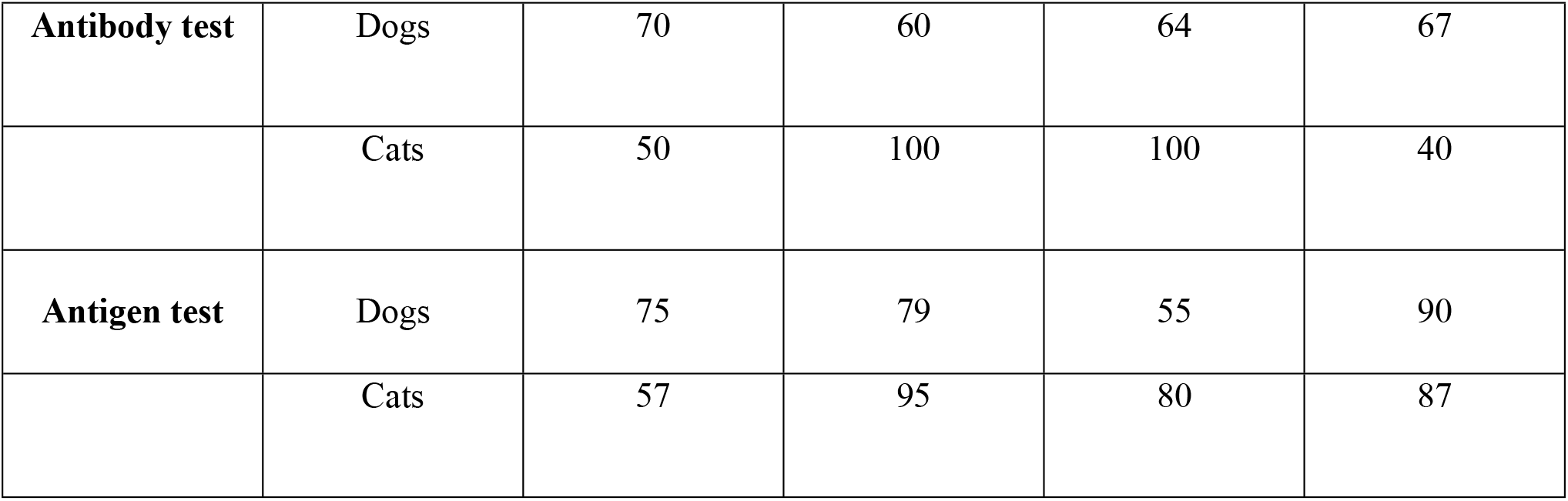
Sensitivity, specificity, positive predictive value (PPV), and negative predictive value (NPV) estimates using human COVID-19 rapid antibody and antigen assays for dog and cat samples, in comparison to PRNT and RT-qPCR methods, respectively.

For dog VTM samples, the RAT was found to have 75% sensitivity and 80% specificity, the PPV was 55% and the NPV was 91%, using the qRT-PCR as the gold standard method. For cat VTM samples, the RAT was found to have 57% sensitivity and 95% specificity, the PPV was 80% and the NPV was 87%.

## Discussion

This study employed human rapid antibody and antigen test kits with dog and cat serum and nasal swab samples, to estimate sensitivity and specificity compared to PRNT and RT-qPCR testing. The test characteristics we observed in dogs and cats are similar to the findings from studies of the same types of tests in humans which have found an average sensitivity and specificity of common commercially available kits in the US range from 50.0-84.3% and 64.5-74.3%, respectively (1,2). Overall, sensitivity of human antibody and antigen tests was moderate in dogs and low in cats, and the specificity was low-to-moderate in dogs and high in cats.

Considering the absence of readily available species-specific rapid SARS-CoV-2 assays for dogs and cats, the test kits in the current study may still be beneficial for field surveillance studies, particularly in low-resource settings.

In our study, all positive animals had positive IgM test bands on the antibody assays suggesting that these animals were recently infected with SARS-CoV-2; this is consistent with the timeframe of animal sample collection occurring during the owners’ acute COVID-19 infections. Previous studies found the opposite; they solely reported the detection of IgG antibodies in animal samples which could mean their samples were collected later in the animals’ infections (51–55). The different results across studies may be due to the variations in collection times from the point of initial exposure of an animal to an infected owner. In the current study, the animal samples were collected while the owners were quarantined at home during an acute COVID-19 infection, supporting the IgM-positive findings. Conversely, other studies collected samples after an owner’s COVID-19 infection had already resolved or the owner’s COVID-19 history was unknown, suggesting that these animals may not have had acute or recent infections. Consequently, the detection of IgM or IgG antibodies in the current or previous studies would align with the animals’ infection stage. –

Several factors could have led to human antigen and antibody SARS-CoV-2 tests exhibiting low sensitivity in dogs and cats, and low specificity in dogs in this study. For example, the test kits were not used in the manner described in the instruction manuals which could have potentially decreased the efficacy of the tests (44,45). Firstly, the COVID-19 antibody test and RAT devices were intended for use in human subjects but instead were used with animal biological samples during this study. Secondly, the standard COVID-19 RAT protocols normally involve collecting a fresh respiratory swab sample (i.e., nasal or throat swab) directly from a human subject, submerging the swab in the buffer solution provided with the test kit, and administering the solution to the test device immediately after collection. Additionally, the nasal and oral swab materials were not collected from animals immediately before administering the tests, which was a major limitation in this study. In the current study, we did not have access to fresh biological samples; instead, we used respiratory swab sample material from animals that were previously collected in VTM, frozen, and thawed prior to administration in the test devices. It is possible that sensitivity could have been increased if the samples were tested immediately after collection from animal subjects.

Congruency between positive RAT and RT-PCR results in respiratory swab samples has been described in previous studies (41,42). This suggests that further investigation employing rapid COVID-19 assays *in vivo* in conjunction with gold standard methods may help determine whether they would produce increased sensitivity and specificity when used with live animal subjects. It is also possible that our storage conditions (i.e., undergoing multiple freeze-thaw cycles) could have resulted in degradation of the SARS-CoV-2 virus and antibodies in the VTM and serum, respectively (56–58). Cats were found to be more susceptible to SARS-CoV-2 infection in multiple studies (59–61). However, it remains unclear whether differences in viral susceptibility between species may have played a role in the higher assay specificity in cats compared to dogs in this study, or if other biological differences between these species influenced viral or antibody detection using these devices. Lastly, there is subjectivity in reading tests and it can be challenging to visually determine whether a test result is positive as the test bands can be very faint, potentially due to low viral load and possibly due to other unknown factors (42).

There is a need for affordable and accessible testing methods for SARS-CoV-2 in domestic dogs and cats to integrate a One Health approach in disease surveillance. The “One Health” concept posits that human, animal, and environmental health are interconnected, with an emphasis on preventing diseases directly or indirectly arising between animals and humans at the human-animal interface (62,63). Considering the proximity of domestic animals and humans, the surveillance methods in the current study could be implemented through a One Health framework to detect disease trends resulting from zoonotic or reverse-zoonotic transmission.

High seroprevalence (21-57%) of SARS-CoV-2 has been found among dogs and cats, with some of the highest rates in COVID-19-positive households, providing evidence of suspected human-to-animal transmission (14,16,18,31,37,42,64–69). It is therefore critical to monitor SARS-CoV-2 and other coronaviruses in susceptible species (e.g., dogs and cats) to mitigate the threat of viral evolution in non-human hosts that could potentially lead to more pathogenic strains that could result in the next pandemic (“Disease X”) (19–26).

In this study of 60 dogs and cats, we found that commonly available human rapid COVID-19 assays could be used to test animal samples and have similar but somewhat lower performance as compared with human samples (1,2). This suggests that commercial RAT and antibody assays could potentially have real-world applications in veterinary settings, surveillance investigations, or households with dogs and cats. It is important to find accessible and affordable testing methods that would allow veterinary and public health professionals to conduct surveillance of SARS-CoV-2 in domesticated dogs and cats because of the potential for cross-species transmission that could negatively affect human or animal health.

## Author contributions

LC conceptualized and designed the study, conducted the experiment, curated the data, performed statistical analysis, wrote the original draft, revised the manuscript, secured funding, and managed project logistics and coordination. SH, GH, and the Hamer Laboratory team at Texas A&M University collected biological samples from animal subjects in the field, tested them using gold-standard RT-PCR and PRNT methods, and provided sera and VTM samples to the partnering investigator in New York. SH also contributed to study conceptualization, design, and manuscript review and editing. CT and GJ assisted with data analysis and manuscript review and editing. JG supervised the research project, contributed to study conceptualization and design, conducted the experiment, secured funding, and participated in manuscript review and editing.

## Acknowledgments

We thank Lisa Auckland and Wendy Tang at Texas A&M University for their significant contributions to this work, including the provision of biological samples and the conduct of RT-PCR and PRNT analyses. We also acknowledge Dr. Catherine Machalaba, Dr. William B. Karesh, and the staff at the City University of New York Graduate School of Public Health and Health Policy for their mentorship and guidance during the development of this manuscript. We are especially grateful to Dr. Xinyin Jiang for permitting us to use her laboratory facilities at Brooklyn College.

## Supporting information

**S1 Table. Comparative diagnostic performance of antibody and antigen testing in canines and felines**. This table summarizes the sensitivity, specificity, positive predictive value (PPV), and negative predictive value (NPV) for both antibody-based and antigen-based diagnostic assays across dog and cat populations.

**S2 Dataset. Diagnostic accuracy of serum testing in veterinary samples**. Summary of diagnostic performance metrics. This file contains the contingency table values (TP, FP, FN, TN) and calculated performance metrics including prevalence, sensitivity, specificity, and predictive values for all samples combined, as well as canine and feline subgroups.

**S3 Dataset. Diagnostic accuracy of VTM testing in veterinary samples**. This table presents the diagnostic performance metrics for VTM-based testing. It includes raw counts for True Positives (TP), False Positives (FP), False Negatives (FN), and True Negatives (TN), alongside calculated values for prevalence, sensitivity, specificity, positive predictive value (PPV), and negative predictive value (NPV) for all samples and species-specific subgroups.

